# Elevated CO_2_ and warming change the nutrient status and use efficiency of *Panicum maximum* Jacq

**DOI:** 10.1101/792440

**Authors:** Juliana Mariano Carvalho, Renato de Mello Prado, Rafael Ferreira Barreto, Eduardo Habermann, Roberto Botelho Ferraz Branco, Carlos Alberto Martinez

## Abstract

*Panicum maximum* Jacq. ‘Mombaça’ (guinea grass) is a C_4_ forage grass widely used in tropical pastures for cattle feeding. In this study, we evaluated the isolated and combined effects of warming and elevated CO_2_ concentration [CO_2_] during summer on the nutrient content, nutrient accumulation, nutrient use efficiency and growth of *P. maximum* under field conditions with adequate water supply. The temperature and [CO_2_] in the field were controlled by temperature free-air controlled enhancement and free-air CO_2_ enrichment systems, respectively. We tested two levels of canopy temperature: ambient temperature and 2°C above ambient temperature, as well as two levels of atmospheric [CO_2_]: ambient [CO_2_] (aCO_2_) and 200 ppm above ambient CO_2_ (eCO_2_). The experiment was established in a completely randomised design with four replications, in a 2×2 factorial scheme. After the pasture establishment, plants were exposed to the treatments for 30 days, with evaluations at 9, 16, 23 and 30 days after the treatments started. Results were dependent on the time of the evaluation, but in the last evaluation (beginning of the grazing), contents of N, K, Mg and S did not change as a function of treatments, P decreased as a function of warming, in [aCO_2_] and [eCO_2_], and Ca increased under [eCO_2_] combined with warming. There was an increase in root dry mass under warming treatment. Combined treatment increased N, Ca and S accumulation without a corresponding increase in the use efficiency of these same nutrients, indicating that the fertiliser dose should increase in the next decades due to human-induced climate change. Our short-term results suggest that the combination of high [CO_2_] and temperature will increase *P. maximum* productivity and that the nutritional requirement for N, Ca and S will increase.

## Introduction

During the last decades, anthropic emissions of greenhouse gases, such as carbon dioxide (CO_2_), nitrous oxide (N_2_O) and methane (CH_4_), have induced alterations in the natural climate cycles of the Earth, elevating the mean surface temperature of the planet [1,2]. The global temperature has been increasing in the last years, and several climate models estimate that this trend will continue in the next decades [3]. Many climate change scenarios have been proposed, depending on the future emissions of greenhouse gases and mitigation policies. According to a moderate-impact scenario outlined by the Intergovernmental Panel on Climate Change (IPCC), the atmospheric CO_2_ concentration ([CO_2_]) will reach 600 ppm by 2100, while the global surface temperature will be between 2.0 and 3.7°C above the pre-industrial average temperature [3].

In tropical and sub-tropical regions, livestock is one of the most important economic activities, and pastures cover extensive areas of the territory, being the main source for cattle feeding in most of these regions [4]. The effects of climate change on the nutritional composition of tropical forage plants deserves attention because climate change factors might alter nutrient uptake and nutrient use efficiency (NUE) by plants, affecting pasture productivity, forage quality and livestock [5].

The responses of tropical plants to elevated [CO_2_] are poorly understood when compared with species grown in temperate and sub-tropical regions, especially C_4_ species. However, some general responses may be highlighted. For example, increased [CO_2_] decreases transpiration rates and increases photosynthesis in many species, thereby greatly increasing water use efficiency [6]. The fact that transpiration governs the root– ion contact of N, Ca, Mg and S [7] suggests that at high [CO_2_], less absorption of these nutrients may occur. Accordingly, it was shown that elevated [CO_2_] led to decreased transpiration and less uptake of N, K, Ca, Mg and S in wheat plants, although the differences were dependent on the time of evaluation. In addition, as the amount of nutrients absorbed per unit of transpired water increased with elevated [CO_2_], the authors indicated that high [CO_2_] is not the only factor responsible for decreased nutrient uptake [8]. However, increased photosynthesis suggests that more nutrients are needed to sustain plant growth, or that NUE is increased. NUE refers to the ability of the plant to convert absorbed nutrient to dry matter [9]. In addition, high [CO_2_] may increase root development and modify the root foraging strategies in order to obtain more resources and sustain higher plant growth [10]. However, an increment of atmospheric [CO_2_] will be followed by an increase in temperature [3].

C_4_ species are adapted to warm, as well as arid environments. Experiments suggest that *P. maximum* would benefit by a 2°C warming under well-watered conditions, by exhibiting increased dry mass [11] and not showing increased stress indicators, such as malondialdehyde and hydrogen peroxide [12]. However, under heating, there was no increase in photosynthesis and transpiration [6,13], suggesting that under a warmed atmosphere, *P. maximum* may exhibit increased NUE. It was also observed that under heating, *P. maximum* exhibited an increase in the concentration of many amino acids, such as valine, threonine and phenylalanine, which have N in their structure [14]. In addition, warming may stimulate nutrient uptake through increased root system growth, increased nutrient diffusion rates and water in-flow [15] but at the same time, gain in dry mass on heating may result in a leaf N dilution effect in *P. maximum* [11]. The same authors observed that the gain in leaf dry mass remains under conditions of combined warming and elevated [CO_2_], but leaf N content increases, suggesting that combined warming with elevated [CO_2_] results in lower N use efficiency. For other macronutrients, no published studies have evaluated the mineral composition and NUE of *P. maximum* under combined effects of elevated [CO_2_] and warming, to date.

Therefore, we exposed a field-grown pasture of guinea grass to elevated [CO_2_] and warming using a combination of temperature free-air controlled enhancement (T-FACE) and free-air CO_2_ enrichment (FACE) systems in order to understand the nutrient dynamics under a short-term experiment. We hypothesise that elevated [CO_2_] may decrease nutrient content but increase nutrient accumulation and NUE, while warming could increase nutrient content and nutrient accumulation but decrease NUE, mainly due to the possible increase in biomass production. In addition, we hypothesised that *P. maximum* dry mass in the warming and elevated [CO_2_] combined treatment will have no difference relative to warming alone, but that nutrient content and accumulation will increase, and NUE will decrease.

## Material and methods

### Study site and system

The study was carried out at the Trop-T-FACE facility, located at the University of São Paulo (USP), Campus of Ribeirão Preto (São Paulo, Brazil), at 21°10’8” S and 47°51’48.2” W and 580 m altitude. The Trop-T-facility is composed of the T-FACE and the FACE systems. According to Thornthwaite [16], the climate at the facility is classified as B2rB’4a’, moist mesothermal with a small water deficiency. The soil was classified as dystrophic red Latosol [17].

Two months before sowing, we collected 10 soil samples (20 cm deep) at the experimental area and analysed it for fertility [18], obtaining the following results (Table 1).

**Table 1.**
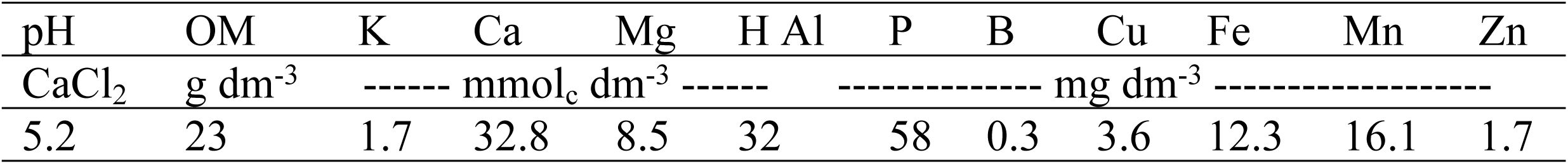
Soil chemical analysis (0 to 20 cm depth) at Trop-T-FACE facility.

Soil preparation consisted of the soil rotation and the application of calcined limestone (48% CaO and 16% MgO) to correct for soil acidity (increasing soil pH to 5.5), using a cultivator 2 months before seeding. Soil fertilisation was conducted [19] through the mechanical incorporation of the fertilisers to 0.10 m depth. The following sources were used: simple superphosphate (18% P_2_O_5_), potassium chloride (60% K_2_O), zinc sulphate (22% Zn), boric acid (17% B) and sodium molybdate (39% Mo). During the experimental period, the accumulated rainfall was 224 mm, and the average temperature was 25 °C, with minimum and maximum of 16 and 35 °C, respectively, while the average air relative humidity was 87% [6].

### Description of treatments and planting method

We tested two canopy temperatures: ambient temperature (aT) and 2°C above ambient temperature (eT) and two atmospheric [CO_2_]: ambient [CO_2_] (aCO_2_) and 200 ppm above aCO_2_ (eCO_2_). The experiment was set up in a completely randomised design, with four replications in a 2×2 factorial scheme. Treatment combinations were designated as follows: aTaCO_2_ (ambient temperature and ambient [CO_2_]), eTaCO_2_ (elevated temperature and ambient [CO_2_]), aTeCO_2_ (ambient temperature and elevated [CO_2_]) and eTeCO_2_ (elevated temperature and elevated [CO_2_]). Seeds of *P. maximum* cv. Mombaça were sown manually in 16 plots (10 m × 10 m), with a final planting density of 16 plant m^-2^ [20]. During seedling growth, supplemental irrigation was performed when necessary. After the pasture establishment, when plants reached 90 cm in height, a standardisation cut was performed at 30 cm above the ground, as part of the post-grazing management. Then, plants were exposed to the treatments for 30 days. In rotational grazing practices, 30 days is the normal plant re-growth time that is often used for this species [21]. Studies indicated that the maximum browsing efficacy of *P. maximum* cv. Mombaça and the highest leaf dry mass production are achieved with 30 cm post-grazing pasture height and 90 cm pre-grazing targets, respectively [22]. In tropical zones, guinea grass is often cultivated under rain-fed conditions, so we decided not to irrigate the plants after the pasture establishment.

Treatments were applied inside circular plots consisting of a 2-m-diameter ring (equivalent to 3.14 m^2^), placed in each 10 m × 10 m plot. We used a safety distance of 12 m between experimental plots to avoid CO_2_ cross-contamination.

### Trop-T-FACE facility description

The eCO_2_ treatment was applied using a FACE system [23]. In each eCO_2_ and eCO_2_ + eT plot, a PVC ring with micro-apertures was used to fumigate pure CO_2_ into the plant canopy. A control unit regulated the amount of CO_2_ required in eCO_2_ and eCO_2_ + eT plots to increase [CO_2_] 200 ppm above the level in the plots with [aCO_2_]. The [CO_2_] in each plot was monitored by a portable [CO_2_] sensor model GTM220 (Vaisala, Finland) located in the centre of each plot, at canopy height. To regulate the opening of the solenoid valves, the central control unit used a proportional integral derivative (PDI). CO_2_ was stored in liquid form in a 12-t cryogenic tank and vaporised before being sent for distribution to the FACE control unit. CO_2_ supply to achieve high-CO_2_ levels was regulated through automatic electromagnetic regulators (ITV model, SMC Corporation, Japan) [23,24]. CO_2_ fumigation by the FACE system occurred daily, from sunrise to sunset.

The eT treatment was applied using a T-FACE system [25]. In each warmed plot, six infrared Salamander heaters (1000 W, 240 V; model FTE-1000, Mor Electric Heating, Comstock Park, MI, USA) were used to warm the plant canopy to 2ºC above the ambient canopy temperature. In each plot, we used an infrared radiometer model SI-1H1-L20 (Apogee Instruments, UT, USA) to monitor the canopy temperature. Using temperature data of warmed and non-warmed plots, the central unit of the T-FACE was calculated, and the voltage of the resistors was adjusted in each heated plot to reach the set-point. Our set-point in this experiment was 2ºC above the ambient canopy temperature.

### Plant growth

At 9, 16, 23 and 30 days after treatment (DAT), plants were collected using the square method, with an area of 0.0625 m^2^, and cut with scissors at 0.30 m above ground. Plant material was washed in running water, neutral detergent (0.1%), HCl (0.3%) and deionised water, and then packed in paper bags and oven-dried at 60°C with forced air circulation until a constant mass was obtained.

To determine the root growth, at 30 DAT, we collected two samples per plot using a soil sampling probe (Sondaterra, Brazil) with an 11-cm internal diameter and volume of 1,900 cm^3^. Samples were collected at a soil depth of 0–0.20 and 0.20–0.40 m, respectively. Each sample was washed, and the roots were dried at 70°C for 72 h to determine the root dry mass.

### Nutrient composition

The above-ground dry mass was milled in a Willey-type mill. N was determined from the digestion of samples in H_2_SO_4_, distillation with the Kjeldahl distiller and titration in H_2_SO_4_ solution [26]. From the digestion of the samples in HNO_3_ and perchloric acid solution, P was determined in a spectrophotometer from the phosphovanadomolybdic complex formed in the reaction of P with the solution of molybdovanadate, while K, Ca and Mg were determined by atomic absorption spectrophotometry, and S was measured by turbidimetric determination of the barium sulphate suspension after the addition of barium chloride [26]. We calculated the shoot nutrient accumulation (NA) based on the shoot nutrient content and the shoot dry mass:

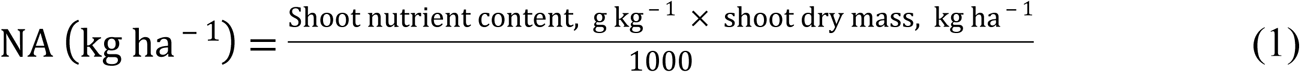

The NUE was calculated according to Siddiqi and Glass [9], expressed as follows:

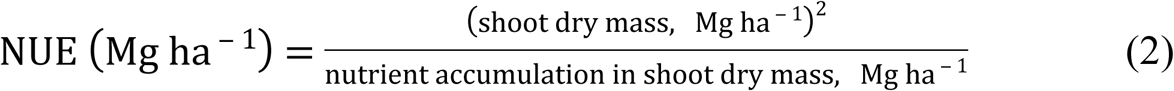

### Statistical analysis

The data normality was checked by the Shapiro–Wilk test. We used a factorial two-way analysis of variance (ANOVA) to test the main effects of [CO_2_] and temperature, as well as their interaction when factors were combined. Non-significant means were compared by the F test. Means of significant interactive effects were compared using Tukey’s test (*p* ≤ 0.05). Analyses were performed using Sisvar software [27]. Trends between nutrient content and shoot dry mass were tested using regression analysis performed in GraphPad Prism software.

## Results

### Macronutrient content

*Panicum maximum* shoot N content was not affected by treatments, except at 16 DAT when plants grown under eT, independently of [CO_2_] level, showed decreased N content (Fig 1A, S1). Differences in P content were observed only in the last sampling (30 DAT) when plants grown under eT, regardless of [CO_2_], showed lower P contents (Fig 1B, S1). In general, plants grown under eTaCO_2_ had lower K content, although [eCO_2_] mitigated this reduction in some samplings (Fig 1C, S1). Differences in Ca content were observed only in the last two evaluations; at 23 DAT, warming, regardless of [CO_2_], resulted in higher Ca content, and at 30 DAT, there was an interaction, so the eTeCO_2_ treatment resulted in higher Ca content than the other treatments (Fig 1D, S1). Mg content increased at 9 DAT under eCO_2_aT treatment (Fig 1E, S1). The S content was not altered by the treatments (Fig 1F, S1).

**Fig 1.**
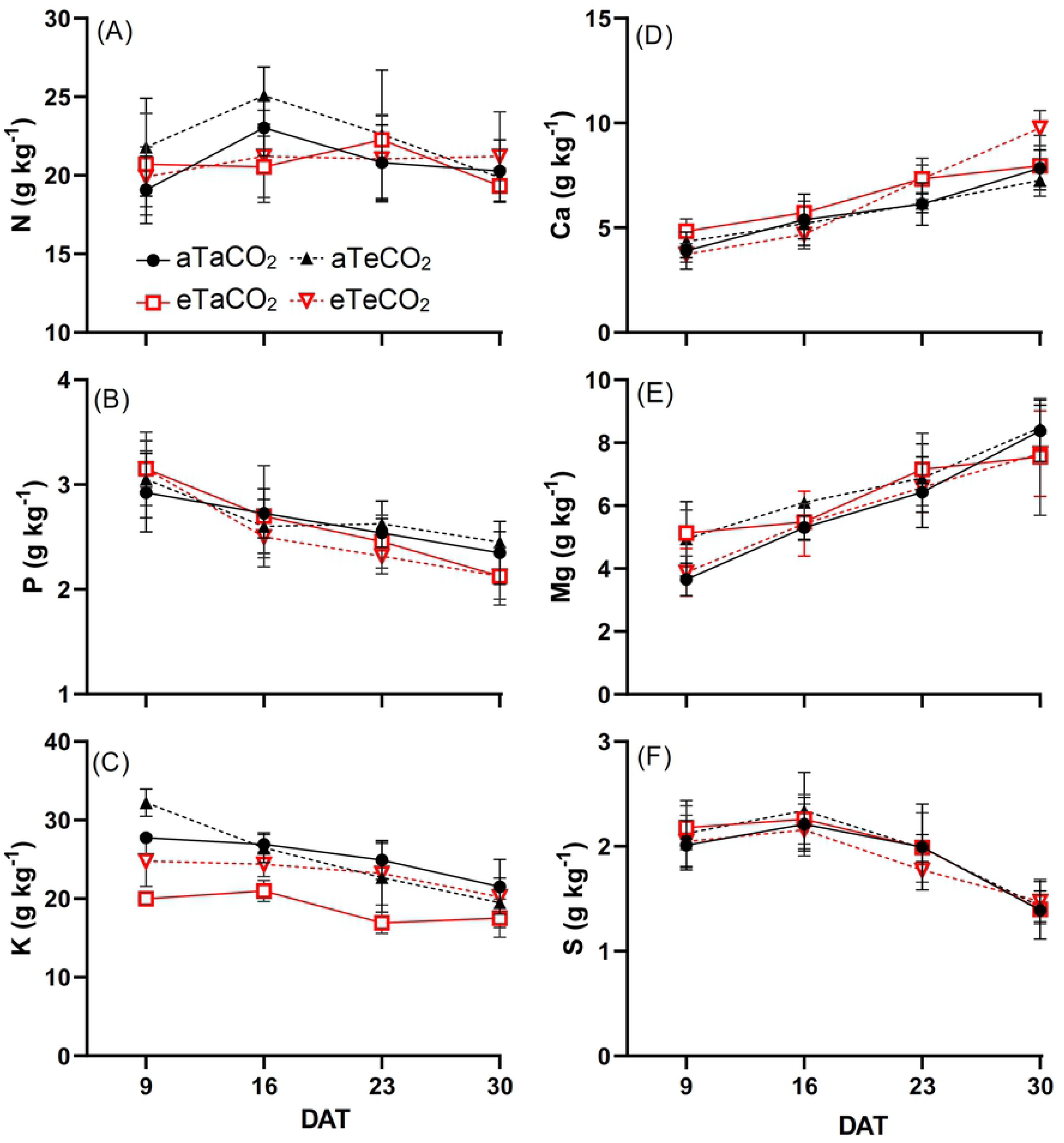
N (A), P (B), K (C), Ca (D), Mg (E) and S (F) content of *P. maximum* shoot during the experiment at 9, 16, 23 and 30 days after treatments (DAT). Treatments: aTaCO_2_ (ambient temperature and ambient [CO_2_]), eTaCO_2_ (2°C above ambient temperature and ambient [CO_2_]), aTeCO_2_ (ambient temperature and 200 ppm above ambient [CO_2_]), and eTeCO_2_ (2°C above ambient temperature and 200 ppm above ambient [CO_2_]). Bars show means and SE of four replicates.

### Macronutrient accumulation

At 16 DAS, [eCO_2_], regardless of temperature level, increased N accumulation; at 23 DAT, eT and [eCO_2_] alone resulted in increased N. At 30 DAT, eT, regardless of [CO_2_] level increased N accumulation (Fig 2A, S1). At 9 DAT, eT, regardless of [CO_2_] level, increased P accumulation, while at 23 and 30 DAT, interactions between the factors increased the P accumulation (Fig 2B, S1). At 16 and 23 DAT, K accumulation increased as a function of [eCO_2_], regardless of the temperature (Fig 2C, S1). At 23 and 30 DAT, Ca accumulation increased as a function of eT, regardless of [CO_2_] level (Fig 2D, S1). At 9 DAT, the interactive effect indicated an increase of Mg accumulation in the eTaCO_2_ treatment. At 16 DAT, plants grown under [eCO_2_] in both temperature conditions had higher Mg accumulation than the other plants. At 23 DAT, there was an interaction again, so that all treatments resulted in an increase of Mg accumulation relative to aTaCO_2_. At the last evaluation, that is, at 30 DAT, there was no effect of treatments on Mg accumulation (Fig 2E, S1). We observed an increase of S accumulation only at 30 DAT under eT plots, regardless of the [CO_2_] level (Fig 2F, S1).

**Fig 2.**
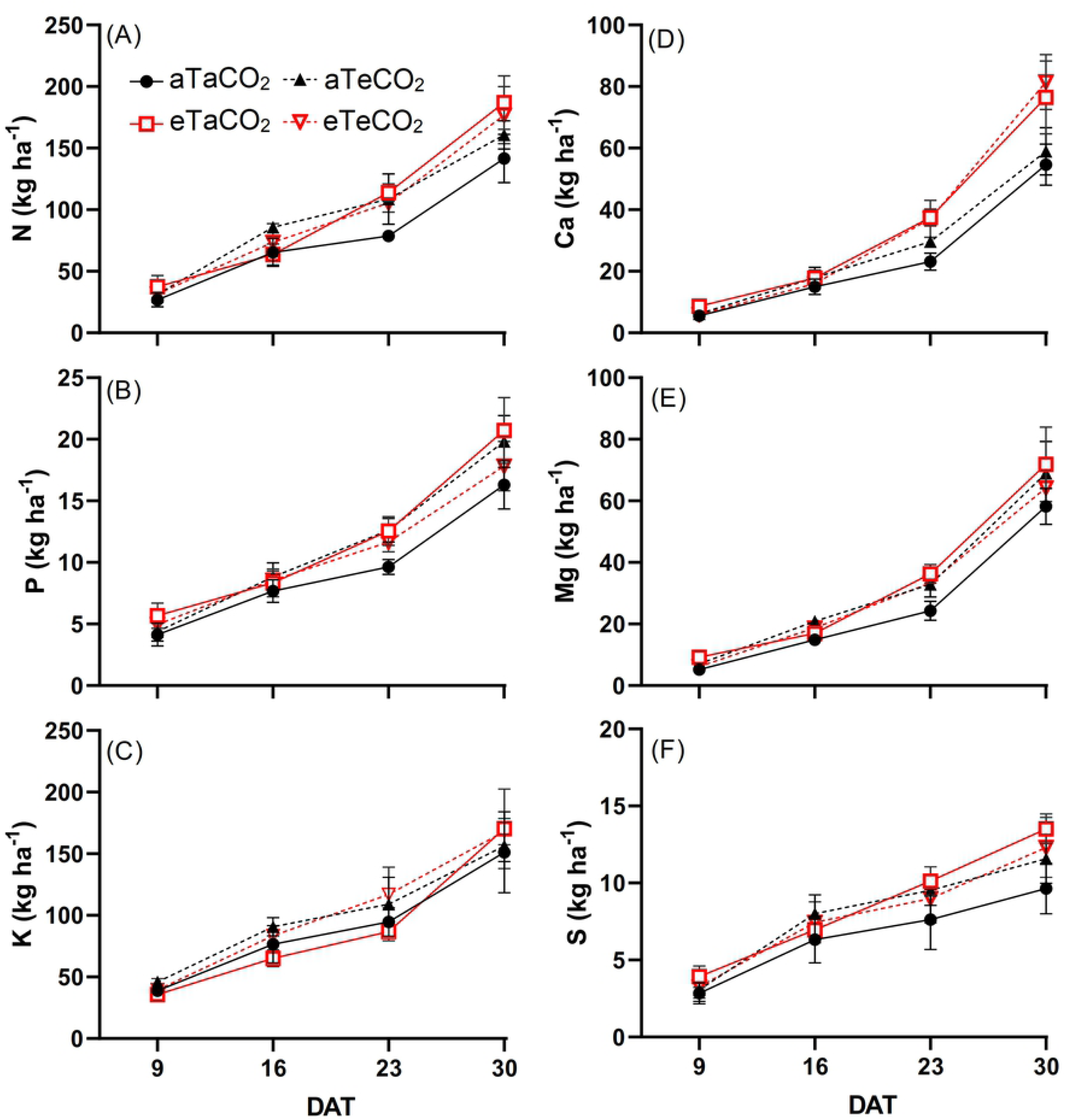
N (A), P (B), K (C), Ca (D), Mg (E) and S (F) accumulation of *P. maximum* shoot during the experiment at 9, 16, 23 and 30 days after treatments (DAT). Treatments: aTaCO_2_ (ambient temperature and ambient [CO_2_]), eTaCO_2_ (2°C above ambient temperature and ambient [CO_2_]), aTeCO_2_ (ambient temperature and 200 ppm above ambient [CO_2_]), and eTeCO_2_ (2°C above ambient temperature and 200 ppm above ambient [CO_2_]). Bars show means and SE of four replicates.

### Macronutrient use efficiency

In three of four evaluation days, eT increased the NUE of N, with [eCO_2_] partially mitigating this increase in the last sampling (Fig 3A, S1). At 23 and 30 DAT, NUE of P increased under eT, regardless of the [CO_2_] (Fig 3B, S1). We observed a noticeable tendency of increased NUE of K under eT plots, with [eCO_2_] amplifying this effect (Fig 3C, S1). NUE of Ca increased at 16 DAT under eCO_2_, regardless of the temperature, while at 30 DAT, there was an interaction between treatments for NUE of Ca and Mg, so that only plants grown under eT and [aCO_2_] had higher use efficiency of these nutrients (Fig 3D and 3E, S1). For NUE of S, we also observed a tendency of increase under eT, regardless of the [CO_2_] level (Fig 3F, S1).

**Fig 3.**
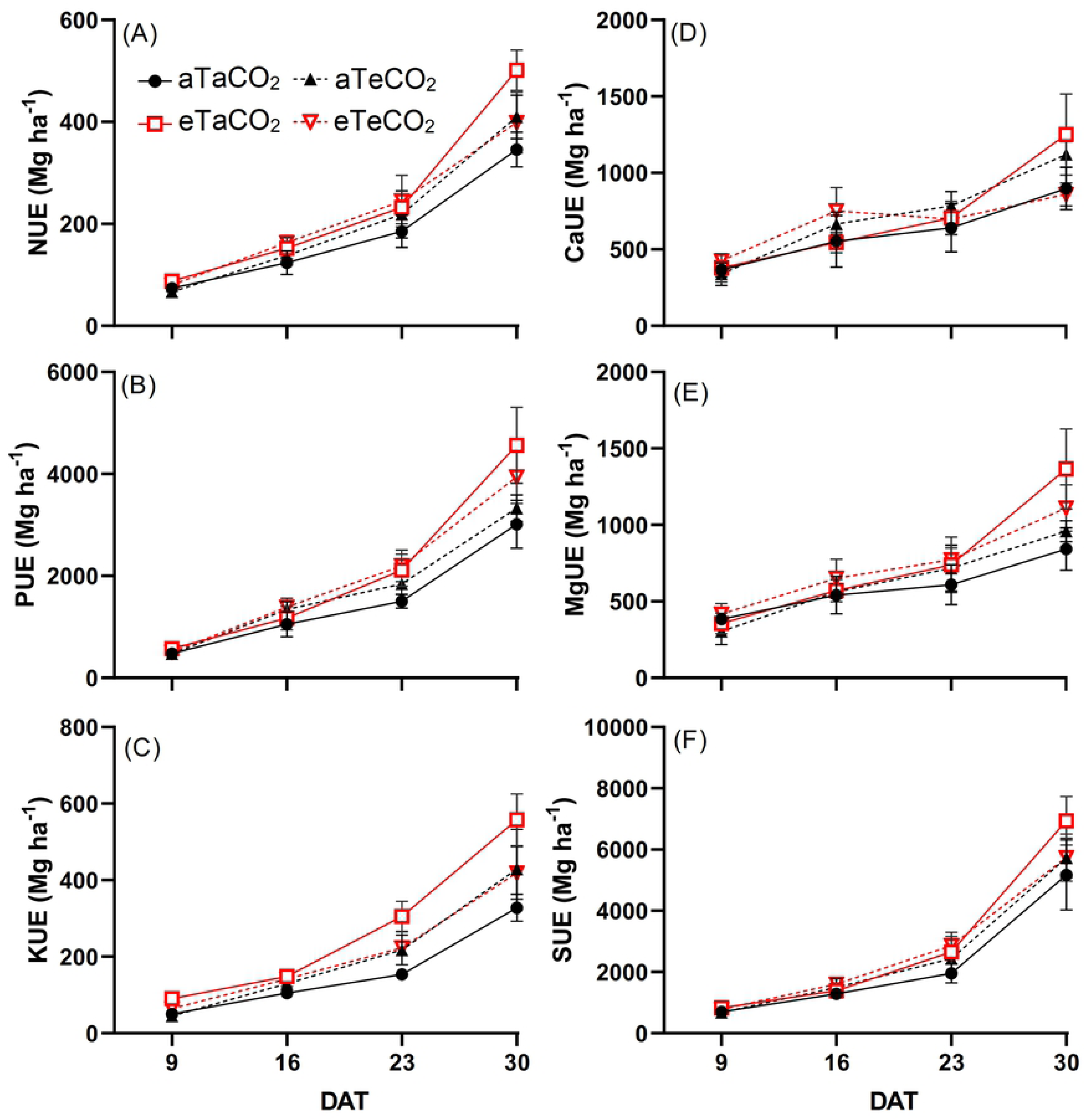
N (A), P (B), K (C), Ca (D), Mg (E) and S (F) use efficiency of *P. maximum* shoot during the experiment at 9, 16, 23 and 30 days after treatments (DAT). Treatments: aTaCO_2_ (ambient temperature and ambient [CO_2_]), eTaCO_2_ (2° C above ambient temperature and ambient [CO_2_]), aTeCO_2_ (ambient temperature and 200 ppm above ambient [CO_2_]), and eTeCO_2_ (2°C above ambient temperature and 200 ppm above ambient [CO_2_]). Bars show means and SE of four replicates.

### Linear regressions between nutrient content and shoot dry mass

N content remained stable, as dry mass increased (Fig 4A). The contents of P, K and S decreased as a function of dry mass increase (Fig 4B, 4C and 4F), suggesting that for these nutrients, there was a dilution effect. Ca and Mg contents increased with increasing shoot dry mass (Fig 4D and 4E).

**Fig 4.**
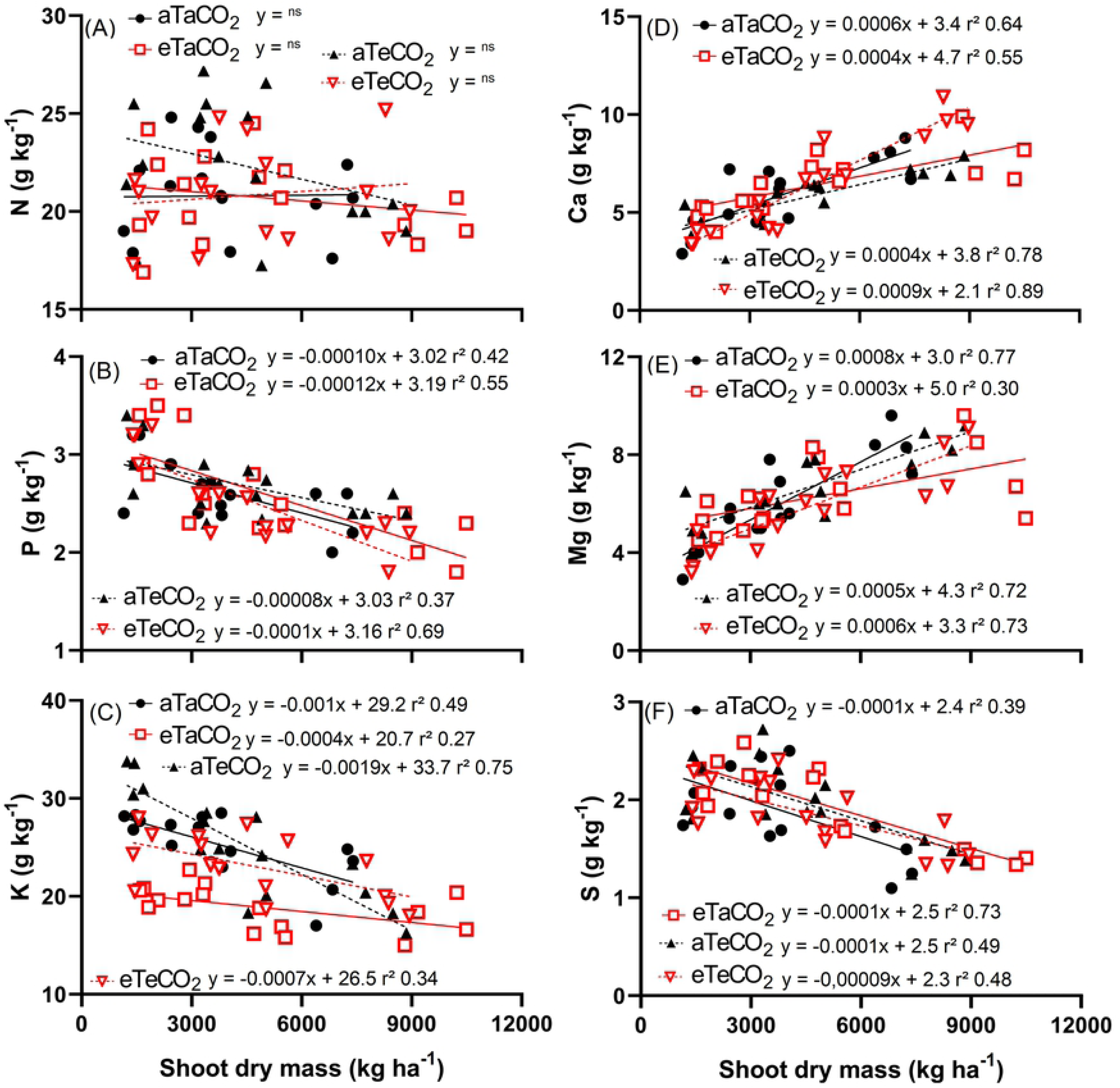
Linear regressions between N (A), P (B), K (C), Ca (D), Mg (E) and S (F) content and shoot dry mass of *P. maximum*. Treatments: aTaCO_2_ (ambient temperature and ambient [CO_2_]), eTaCO_2_ (2°C above ambient temperature and ambient [CO_2_]), aTeCO_2_ (ambient temperature and 200 ppm above ambient [CO_2_]), and eTeCO_2_ (2°C above ambient temperature and 200 ppm above ambient [CO_2_]).

### Dry mass

The eT, in 9 days, in both [CO_2_], resulted in an increase in the shoot dry mass. However, at 16 DAT, plants grown under [eCO_2_] had increased dry mass; At 23 DAT, the [eCO_2_], combined or isolated from eT, resulted in an increase in the dry mass of the shoot. At 30 DAT, the highest dry mass was observed only in plants grown under eT and [aCO_2_] (Fig 5, S1). The root dry mass in the 0–20 and 20–40 cm layers were higher due to the increase in temperature (Fig 6A and 6B).

**Fig 5.**
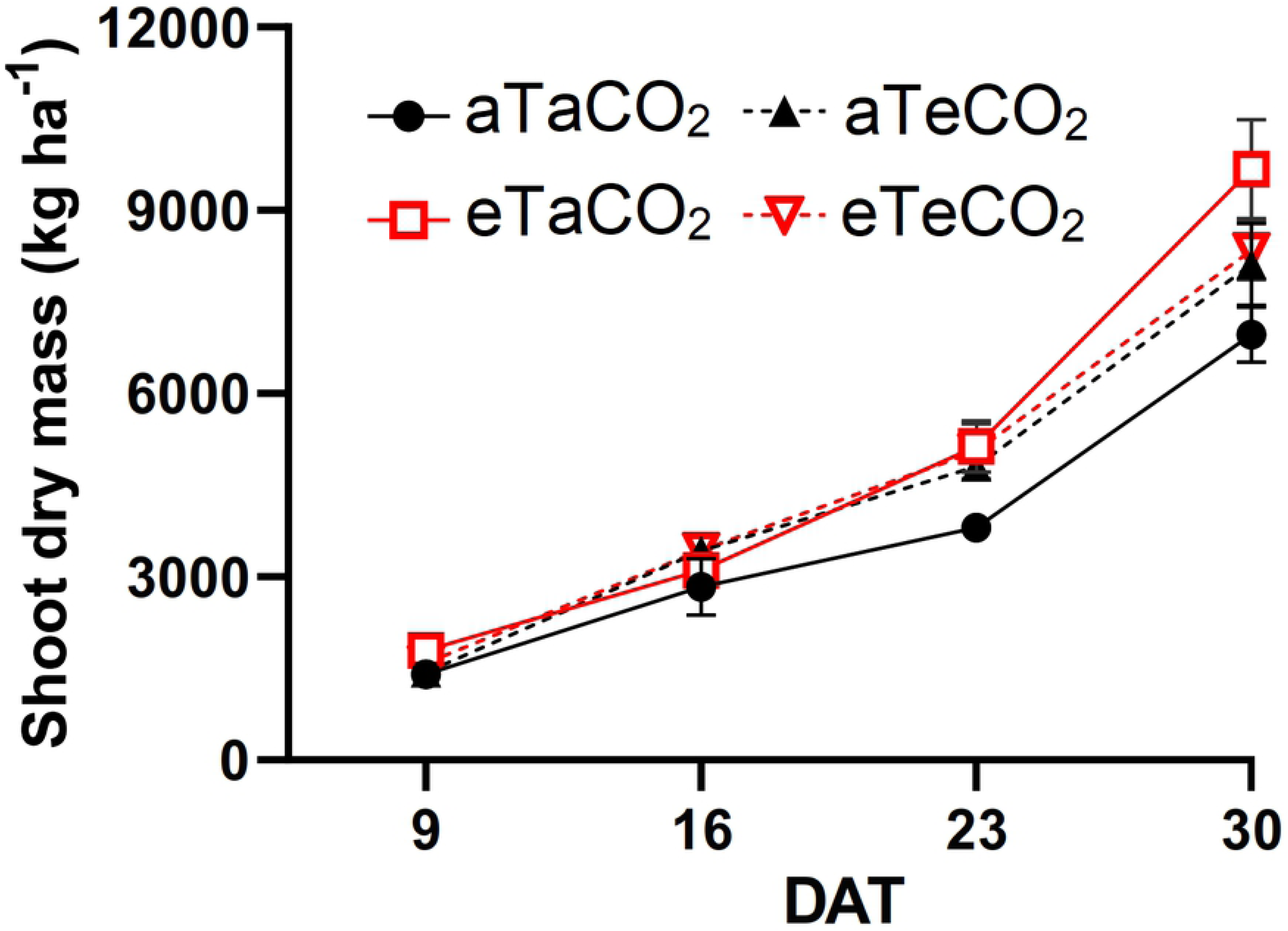
Shoot dry mass of *P. maximum* during the experiment at 9, 16, 23 and 30 days after treatments (DAT). Treatments: aTaCO_2_ (ambient temperature and ambient [CO_2_]), eTaCO_2_ (2°C above ambient temperature and ambient [CO_2_]), aTeCO_2_ (ambient temperature and 200 ppm above ambient [CO_2_]), and eTeCO_2_ (2°C above ambient temperature and 200 ppm above ambient [CO_2_]). Bars show means and SE of four replicates.

**Fig 6.**
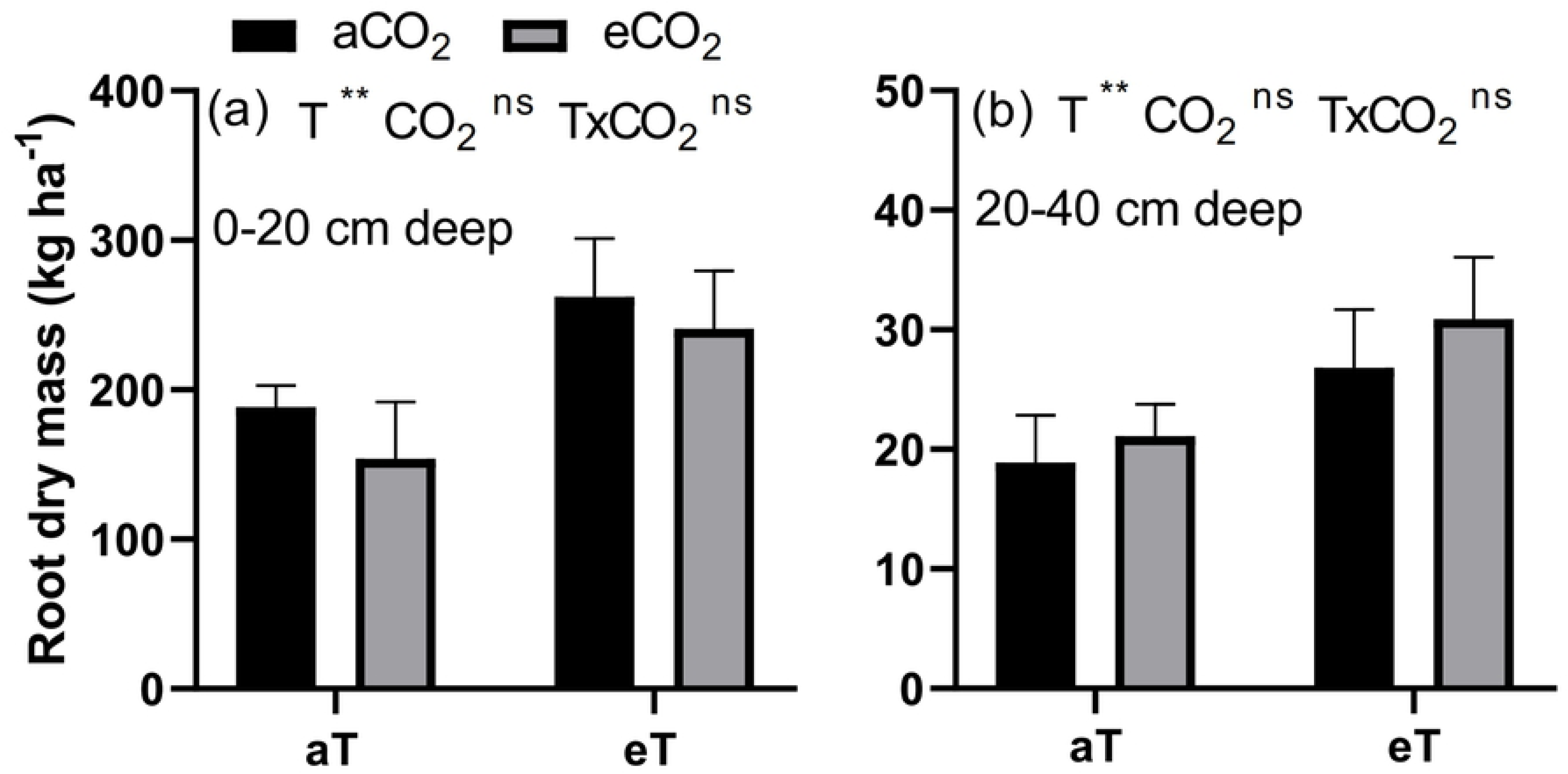
Root dry mass at 0-20 cm deep (A) and at 20-40 cm deep (B) of *P. maximum* at 30 days after treatments. Treatments: aTaCO_2_ (ambient temperature and ambient [CO_2_]), eTaCO_2_ (2° C above ambient temperature and ambient [CO_2_]), aTeCO_2_ (ambient temperature and 200 ppm above ambient [CO_2_]), and eTeCO_2_ (2°C above ambient temperature and 200 ppm above ambient [CO_2_]). Bars show means and SE of four replicates. ^ns^ and ^**^: not significant and significant at 1%, respectively.

## Discussion

Our main hypothesis was not corroborated because the nutrient content, accumulation and NUE responses to eCO_2_ and eT were variable for each nutrient. However, we highlight two major trends observed in our study: i) eT under aCO_2_ promoted an enhanced nutrient accumulation and NUE for most of the nutrients studied and, ii) eCO_2_ had an interactive effect when combined with eT, decreasing the NUE of N, K and Ca. In previous experiments conducted during winter (10–30°C) with adequate water availability, warming combined or not with elevated [CO_2_] resulted in a higher leaf dry mass of *P. maximum*. In addition, the authors observed increased N content in elevated [CO_2_] combined or not with warming, and decreased N content due to warming alone [11]. Here, we find that during summer (16– 35°C), shoot dry mass also increases under eT and [eCO_2_], but the N content was not changed in most evaluations. Moreover, considering the final re-growth phase, which would be the period when the cattle would eat the pasture [21], K, Mg and S were not affected by treatments.

The increased NUE of N under warmed plots with unchanged N content indicated that *P. maximum* acclimates to moderate heating. This response may be associated with changes in N uptake and assimilation, allocation and remobilisation or metabolic modifications [28]. Increased NUE of N may also be achieved by higher transpiration rates. However, recent evidence indicated that a 2°C elevation does not affect transpiration rates or water use efficiency of guinea grass [6]. In addition, we observed no dilution effect as the plants grew. Interesting, this improved NUE of N was detected even under rain-fed conditions, demonstrating that the positive effects of warming demonstrated by other studies under well-watered conditions on *P. maximum* are maintained under rain-fed conditions.

Typically, plants grown under increased temperature exhibit lower P [29]. Accordingly, in our experiment, the decrease in P content, and its higher accumulation under eT, suggest that the dry mass of plants increased with warming and [eCO_2_] and that the P decrease was not limiting for plant growth. In addition, the P use efficiency indicates that the decrease of P content with warming was not harmful to plants. However, the P dilution effect that occurred with increasing plant dry mass does not seem to have been influenced by the [CO_2_] and warming treatments, given the similar slope values of the lines (Fig 4B). However, the increase in photosynthetic rate under elevated [CO_2_] [6] can be associated with the Rubisco content. High [CO_2_] is expected to increase the Rubisco concentration, and this will require more inorganic P to be transformed into organic P for Rubisco synthesis, which is an important component of rRNA involved in enzyme synthesis. Thus, the P use efficiency would increase because a higher proportion of P in plant tissue is used for photosynthesis-associated metabolism and assimilation [30], as we observed here.

In the literature, it is indicated that warming decreased K content in several plant species, although the causes are still unknown [31]. In our experiment, the K content was lower as a result of the temperature increase in the first three evaluations and, in the last one, there was no difference from the other treatments (Fig 1C). Although the K content was lower in the first three evaluations due to eT effects, plants showed increased NUE of K (Fig 3C), which presumably contributed to the gain in dry mass.

The increased Ca accumulation under eT, mainly in the last evaluations (Fig 2D), was probably associated with the increased Ca content (Fig 1D) and the increased shoot dry mass (Fig 5), and also due to no dilution effect (Fig 4D). Moreover, the increased Ca use efficiency (Fig 3D) under this same condition suggests that the gain in dry mass was greater than the increase in its absorption. The increase of Ca accumulation under eT may also be associated with the increased of root dry mass in this same treatment because root interception significantly contributes to Ca uptake [7].

The values close to the Mg and S contents suggest that the changes in the accumulation and the efficiency of these nutrients occurred due to the differences observed in the shoot dry mass and also the fact that although Mg concentrates and S dilutes with increased in shoot dry mass, this does not appear to be influenced by [CO_2_] and warming treatments, due to the similar straight slope (Fig 4E and 4F).

After a long time, the response of C_4_ species to increased levels of [CO_2_] was considered inexistent due to the natural concentration mechanisms of CO_2_ inside the bundle sheath cells. However, there is variation in the [CO_2_] saturation level of C_4_ leaves. While some species appear to be saturated under actual [CO_2_], others are not necessarily saturated at this level [32]. Likewise, *P. maximum* exhibited an increase in photosynthesis as a function of [eCO_2_] [6] and an increase in dry mass, although this result was dependent on the time of evaluation, in our experiment. The increase in shoot dry mass with heating is probably associated with higher efficiency of N, P, K, Ca and Mg use under this same condition (Fig 3). It means that with warming, plants are better able to convert these nutrients into dry mass.

Finally, because the combination of increased [CO_2_] and rising temperatures occur simultaneously [3], our short-term combined treatment results suggest that plants will have decreased P content and increased Ca content. This information may contribute to the interpretation of nutritional diagnoses in the future. Moreover, the increased accumulation of N, Ca and S in the combined treatment, without a corresponding increase in the use efficiency of these same nutrients, indicates that the fertiliser dose may need to be increased in the climate change scenario. Among these macronutrients, decreased efficiency of N use has been reported for other grasses [33].

## Conclusion

Our short-term results suggest that under the combination of [eCO_2_] and eT conditions, *P. maximum* productivity will increase and the nutritional requirement for N, Ca and S will also increase.

## Acknowledgements

We thank Bruce Kimball from the USDA and Franco Miglietta from IBIMET, Italy. The authors thank Wolf Seeds from Ribeirão Preto, São Paulo State, Brazil, for providing seeds of *P. maximum*.

## Supporting information

**Supplementary material 1 (S1).** ANOVA results of macronutrient content, macronutrient accumulation, macronutrient use efficiency, linear regressions, dry mass and all data underlying the findings.

